# How brain operation emerges from its nested recurrent circuitry extending into the body

**DOI:** 10.64898/2026.07.31.742003

**Authors:** Szilvia Szeier, Henrik Jörntell

## Abstract

Behavior and thoughts are dynamic processes driven by activity in a multitude of internal and external nested neuronal circuitry loops. This causes an unexplained complexity in neuron population dynamics. Pursuing an atomic principle of how brain operation emerges from these nested closed loops, here we show that the activity state space travel of a neuron population can be understood as a consequence of a vector field with an attractor point, which emerges from the intrinsic neuronal properties and the synaptic connections between them. Neuronal activity in the brain *in vivo*, however, indicates that attractor point continually moves along trajectories, which we show requires dynamic input from sensors and input-independent activity generation in neurons. In a spinal network-biomechanical system, the state-dependent intrinsic dynamical structures of the neural and the biomechanics systems drive each other’s activity trajectories to generate diverse behaviors. Similarly, independent self-generated thalamic neuron activity forms closed-loop thalamocortical subnetworks that continually shape each other’s trajectories to drive the dynamic global cortical activity also without dynamic sensory input. These high-dimensional attractor trajectories are susceptible to criticality effects that allow for a rich behavioral flexibility based on sensory inputs and intrinsic anticipative subnetworks, explaining why complex neural dynamics is foundational to brain operation. We argue that in this conceptual framework the contribution of each functional subsystem of the CNS to sensorimotor control and motor planning is given a natural, emergent, and scalable explanation, and the goal of synaptic weight learning partially becomes fundamentally different from most models in the literature.

**AUTHOR SUMMARY:** Our paper is inspired by the fact that the understanding of the brain, how it works as an integrated organ, is still poor. We address the question by illustrating the atomic principles of how brain operation emerges from its nested closed loop interactions, internally and through the body. Our focus on emergence and core mechanistic network principles across the CNS provides an inevitability and scalability, which contrasts with the leading neuroscience literature that is fundamentally based on more arbitrary correlations for specific, open-ended assumptions. The emergent dynamical system view that we present flips the table on many aspects of how to interpret brain operation and, not the least, the meaning of learning and how little it can take to transition into adapted behavior.

## INTRODUCTION

The primary function of the brain is to generate diverse behaviors that can increase the chances of survival within its environment ^1^. Sensory information provides the brain with information about the external world in interaction with the body, playing a critical role for continually evaluating the consequences of specific behavioral outputs. In doing so, sensory information closes the loop between the brain, the body, and the environment. Supporting this continuous interaction is the extensive nested and parallel closed-loop, or recurrent, connectivity that also characterizes operation within the central nervous system (CNS) (overview provided in ^2^), giving rise to a system of interacting subnetworks. Most of these subnetworks below the cortical level are primarily reactive ^1^, relying on external sensory inputs to drive their output activity, which in turn impacts that sensory input. In contrast, thalamic neurons can generate activity partly independently of synaptic input ^3–7^, allowing self-sustained dynamic activity in the thalamocortical system. Together with learning and other adaptive processes occurring within individual neurons ^8,9^, these recurrent interactions produce a highly complex dynamical system. In complex systems, collective behavior emerges from the nature of the interactions among the constituent elements and their coupling to the external environment.

Previous work has applied mathematical frameworks to identify structures in neural population activity data ^10,11^ and reported various low-dimensional structures within high-dimensional neural activity patterns ^12^. While these approaches have provided powerful descriptions of population-level neural dynamics, an important open question is how they give rise to functionally meaningful circuitry operation. Ultimately, such descriptions must account for how neural activity generates behavior through continuous interactions among the various subsystems and behavioral subfunctions ^13^ of the nervous system, the body, and the complex environment, to achieve future states supporting survival ^14^. Addressing this challenge may require a complementary perspective that explicitly considers the functional impacts of nested dynamical interactions linking neural activity, sensory feedback, biomechanics, and behavior. Within such a system, the functional contribution of any individual neuron is not fixed but instead depends on the current network state, behavioral context, and ongoing interactions with the environment. Understanding how population-level neural dynamics emerge from these nested interactions is therefore essential not only to understand the fundaments of brain operation, but also for relating recorded neural activity to integrated brain function.

Population level dynamics within the brain are governed by the synaptic connections and the neuronal input-output functions, which can be compactly represented within a vector field framework ^15^. Here we extend this vector field framework to account for the dynamical control of multifunctional behavior in nested recurrent network interactions. To focus on the fundamental emerging principles, we use a minimalistic model of corticospinal behavioral control of biomimetic muscles with biomechanically dependent sensory feedback. We show how independently generated time-varying activity from cortex results in a cascade of continuously changing vector field structures in a spinal interneuron network in closed-loop with the biomechanics, thereby defining continual, non-stationary attractor trajectories throughout the system. Variability from both intended and unintended sources modulates these attractor trajectories, giving rise to within-behavioral variability, as observed in natural movements ^16–18^ and in the underlying neural activity ^19,20^. Importantly, we also find that when such variability is coordinated across sensors and neurons, it can instead drive consistent transitions between distinct attractor trajectories, corresponding to switching behavior, or network solutions ^21,22^. We further discuss how this framework provides a conceptual base to interpret the various CNS components, in terms of how they shape and leverage coupled brain-body-environment dynamics to learn to generate and anticipatively control behavior.

## RESULTS

### Continuously changing cortical activity *in vivo*

Previous work has reported both stationary ^23,24^ and non-stationary (meta-stable) ^25^ dynamics in cortical activity. We determined which regime dominated *in vivo* by examining the time-evolving activity of cortical neurons without applying any patterning filter from embedding methods or conditioning on behavioral parameterizations. We observed that populations of neighboring cortical neurons exhibited complex patterns of continuously changing activity distributions (Fig. 1A-C; fig. S1), here referred to as trajectories within the population activity state space.

**Fig. 1.**
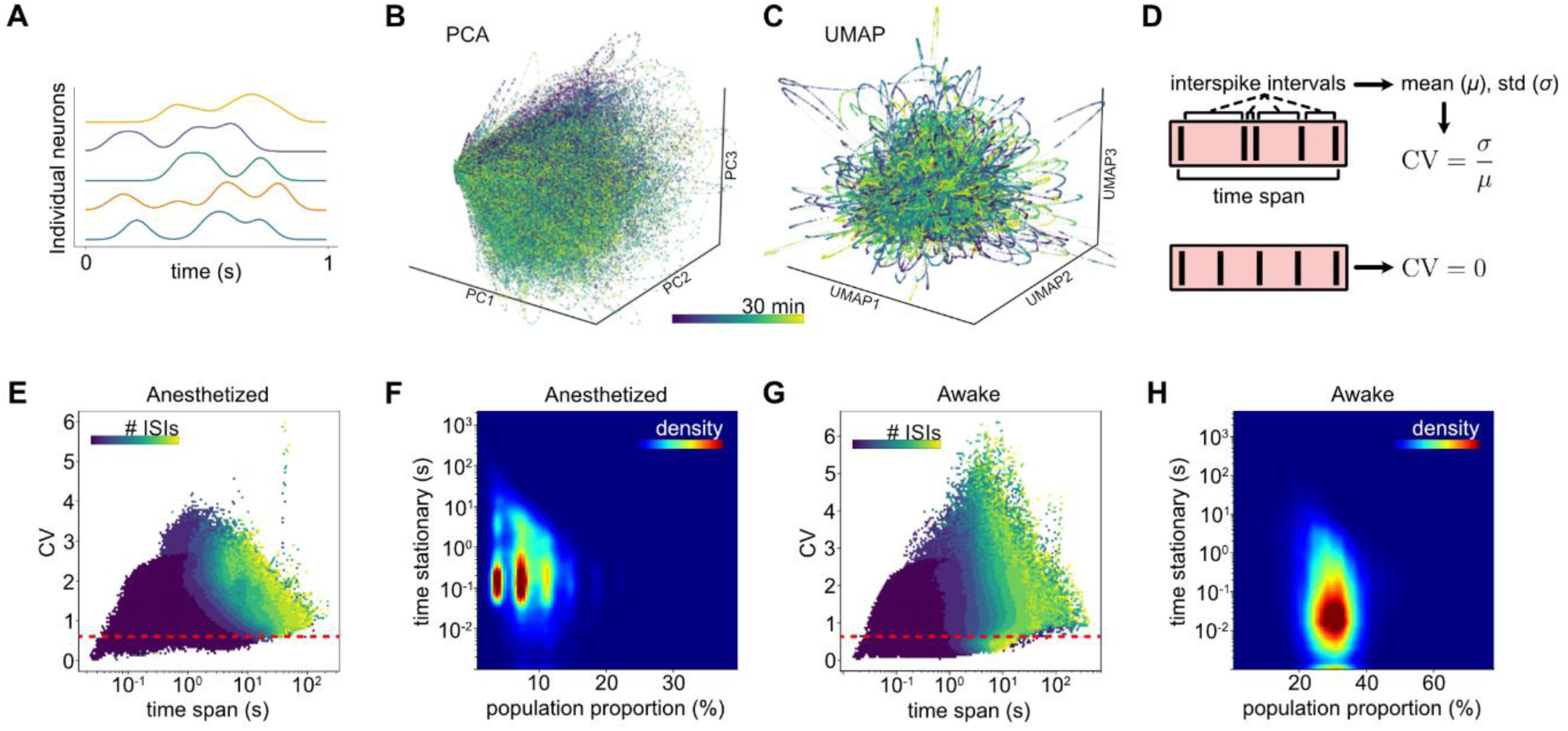
In vivo cortical recordings reveal complex, dynamically changing population activity. A-F) Recordings were obtained from Neuropixels multielectrode array recordings in the primary somatosensory (S1) or primary visual (V1) of the rat neocortex. **(A)** The convolved spiking activity of five neurons during an example interval exhibit continually changing decorrelation patterns in the neuron population activity. **(B)** These changing activity patterns are captured using the first three components of a linear dimensionality reduction (PCA) and; **(C)** a nonlinear approach (UMAP). **(D)** The stationarity of individual neurons was assessed using sequences of consecutive interspike intervals (ISIs). For each sequence, stationarity was quantified by the coefficient of variation (CV, standard deviation divided by the mean) of the ISIs. A higher CV indicates greater variability and zero indicates no variability. **(E)** The relationship between ISI duration and CV for sequences across experiments is colored by the number of consecutive ISIs included in the calculation. Individual neurons consistently showed high variability across interval durations, with progressively larger values as the number of consecutive ISIs increased. **(F)** Population level stationarity, defined as periods during which the stationary intervals (CV<0.6 threshold (red line in e) of multiple neurons overlapped. **(G-H)** Same as in (E-F), but for neural population activity recorded from an awake, head-fixed mice from the International brain laboratory (IBL) dataset ^57^.

To quantify stationarity, we used the coefficient of variation (CV) as a measure of firing variability. For individual neurons, the CV of interspike interval sequences (Fig. 1D) generally indicated high spike firing variability in both anesthetized (Fig. 1E) and awake recordings (Fig. 1G). We used a threshold of CV < 0.6^26^ for stationarity (red line in Fig. 1E, G) to measure the fraction of the neuron population that could be classified as stationary within overlapping time windows over the course of the 1-2 hour recordings (Fig. 1F, H). Such episodes of population-level stationarity were rare (Fig. 1F, H). Across the recordings, the proportion of neurons simultaneously exhibiting stationary activity never exceeded 50%. In general, it was limited to around 7% in anesthetized recordings and 35% in awake recordings, and even then only for brief episodes of 0.1-1 seconds. The lower apparent stationarity under anesthesia could be ascribed to more mechanically stable recording conditions and therefore greater reliability of the unit identification/tracking, as well as the more prominent activity generation in thalamic neurons during sleep. The near absence of sustained widespread stationarity, even in these recordings from spatially confined cortical populations, suggests that stationary regimes are unlikely to dominate across the neuron population at any time. As stationarity basically requires all neurons to participate, we conclude from Fig. 1 that the cortical neuron population has fundamentally non-stationary behavior.

### Vector field properties that emerge from the network itself

To investigate the origin of the continuously evolving patterns of neuron population activity observed *in vivo*, we began by considering the simplest possible closed-loop network (Fig. 2A), a connectivity motif that is ubiquitous in cortical circuits (reviewed in ^2^) as well as through thalamocortical loops ^27^. Neurons in closed loops mutually influence each other’s activity and such networks composed solely of excitatory neurons form positive feedback loops, leading to progressively increasing activity levels (Fig. 2A). In contrast, the addition of inhibitory neurons counteracts this amplification within the closed loop. The full range of these general neuron population-level interactions can be understood through the vector field analogy ^15^, which is an emergent effect from how the neurons mutually influence each other through the network’s connectivity structure and synaptic weights, as well as the activation functions of those neurons.

**Fig. 2.**
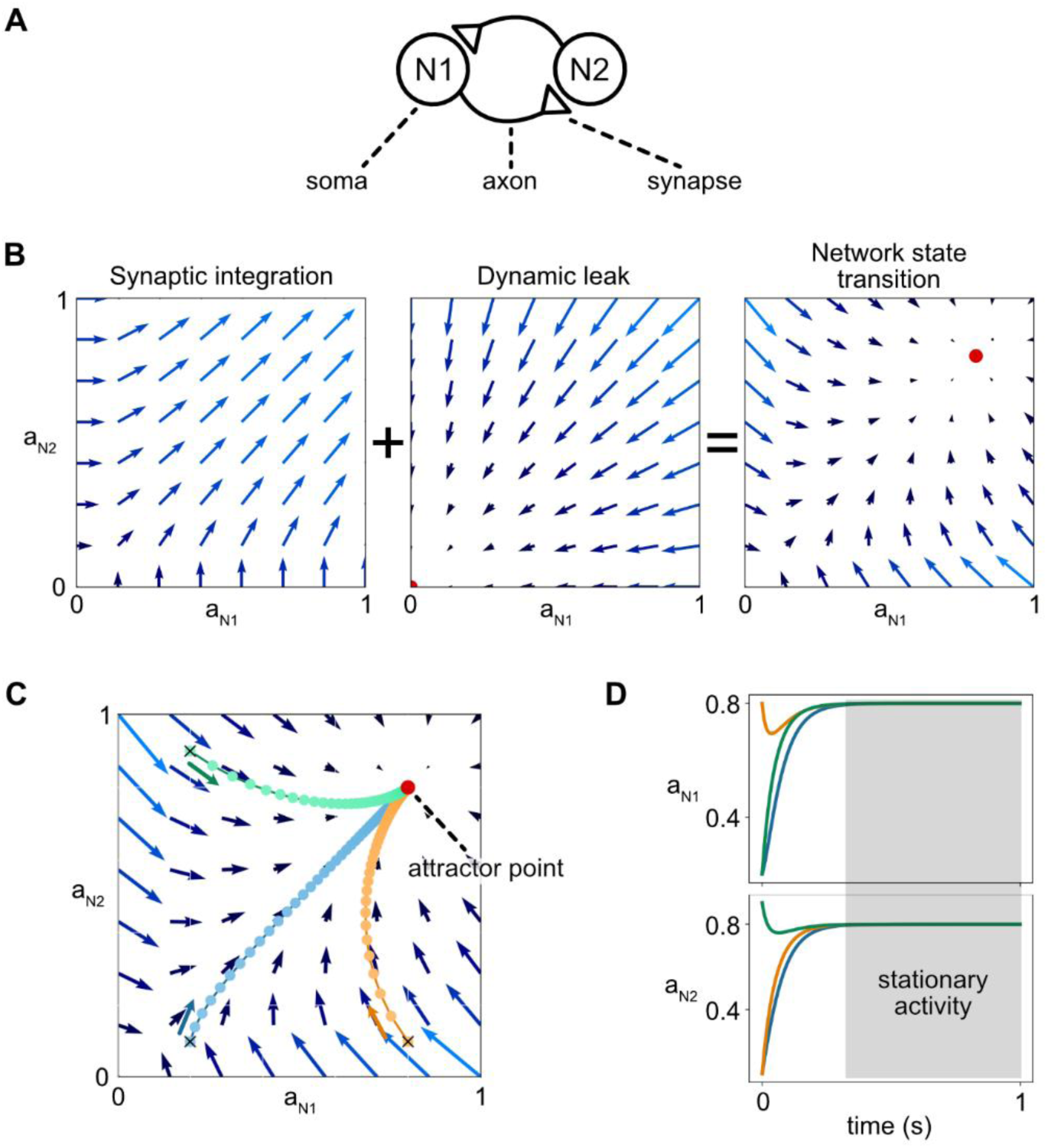
Vector field representation of a recurrent (closed-loop) neuronal network. **(A)** A simple 2-excitatory-neuron network used to explain the principles. **(B)** The vector fields defined by the network synaptic weights ^15^ and the dynamic leak produce a network state transition vector field. **(C)** This vector field has a non-zero attractor point towards which the state of the network converges whenever the activity of at least one neuron is non-zero. **(D)** After a transient convergence phase, the example network remains at its attractor point indefinitely, resulting in stationary activity.

Here, we extend the vector field framework by incorporating a fundamental property of neurons: the membrane acts as a capacitor that integrates the charging caused by synaptic inputs while it also continuously dissipates that charge through its membrane leak channels. This endows neurons with a leaky state memory ^28^, characterized by the membrane time constant, which is here represented by the dynamic leak vector field (Fig. 2B). The combined effects of recurrent excitation and dynamic leak inevitably give rise to a stable state within the vector field, which we term the attractor point. In the fully excitatory network considered here, this attractor point is located at a positive activity level (red dot in Fig. 2C). This effect will emerge wherever a network, or a subnetwork within a network (see below), contains a closed excitatory loop whose overall excitation exceeds its inhibition, generally regardless of the exact connectivity pattern or the latency times within the loop (Fig. S2). In addition, each such network will also have an attractor point at 0 activity.

Under these conditions, regardless of the network’s initial state, the network activity trajectories will converge toward the attractor point (Fig. 2D), where they will remain indefinitely. Consequently, the activity of all constituent neurons becomes stationary. The same behavior occurs in spiking networks, where the position of the attractor in the activity state space corresponds to the resulting average firing rate of the network. This behavior stands in contrast to neuronal population activity observed in the brain, which is persistently non-stationary (Fig. 1). What then enables neurons in closed loop systems to continuously change their activity? Put differently, what mechanism causes the attractor point of the network to continuously remain in motion?

### Activity that is independent of the vector field generates attractor trajectories

Since the synaptic weights that determine the vector field structure of the closed-loop network (Fig. 2A) remain constant over timescales relevant for isolated behaviors ^15^, continual non-repetitive dynamic changes in network activity (Fig. 1) that are in addition robust to longer-term learning effects, require influences originating from outside the vector field. In the brain, two main sources can provide such independent inputs: (1) sensory synaptic inputs and (2) intrinsic active conductances which enable neurons to generate output independently of synaptic input ^3,7,29^. Input-independent activity occurs, for example, in thalamocortical neurons, where hyperpolarization-evoked rebound excitation can generate self-sustained dynamic activity ^6^, i.e. these conductances generate an independent activity state space within the neuron itself. To investigate the effects of such inputs, we added two input channels to the same network used previously. These can be viewed as representing either sensory input or thalamically driven input (Fig. 3A) where the inhibitory input represents an indirect influence mediated through an inhibitory interneuron.

**Fig. 3.**
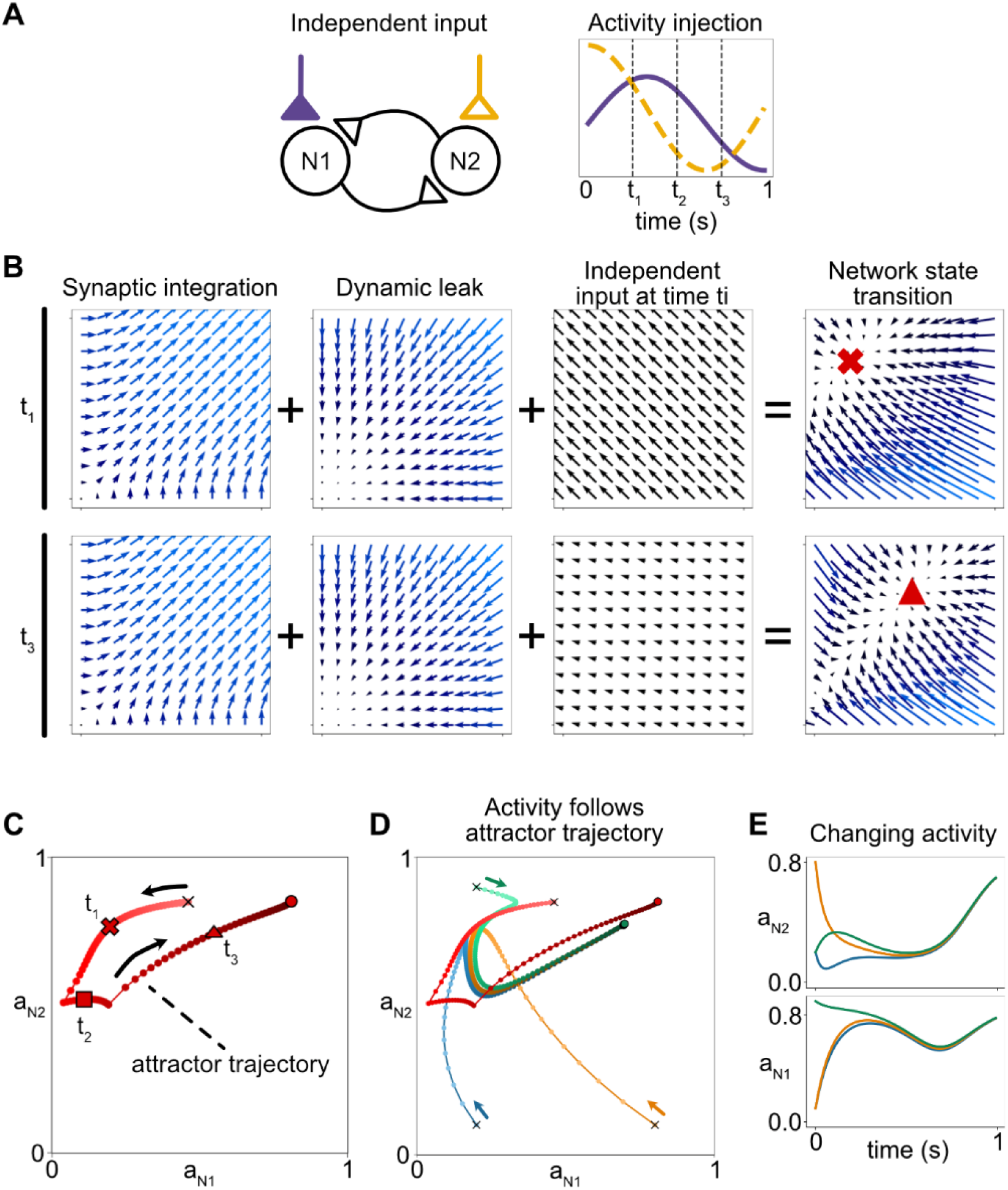
Input creates attractor trajectories and thereby dynamically changing activity patterns. **(A)** The simple network receiving inputs from two independent sources. **(B)** Although the synaptic input and dynamic leak vector fields do not change, the input constitutes an additional vector field that changes between consecutive time steps. The input alters the structure of the vector field and results in a corresponding shift in the attractor point location. **(C)** A sequence of attractor points forms an attractor trajectory. **(D)** The network activity converges towards the attractor trajectory regardless of initial value. **(E)** The attractor trajectory results in dynamically changing neuron activities but still with strong correlations between the neurons.

The addition of these inputs caused the network attractor to move, as illustrated in Fig. 3B for two different phases of the time-varying input. This occurred because the external inputs contributed a third vector field, which combined with the synaptic and dynamic leak vector fields to generate a modified state-transition vector field. As the external input changes over time, so does its impact on the overall vector field, which hence gradually changes structure. Consequently, the attractor moves continuously through the state space, tracing a trajectory over the time course of the input (Fig. 3C).

As in Fig. 2, the network activity is drawn toward the attractor regardless of its initial state. However, because the attractor itself is now moving, the activity strives towards an attractor trajectory rather than converging to a fixed attractor point (Fig. 3D). This results in continuously evolving neuronal activity (Fig. 3E), producing non-stationary network dynamics. Nevertheless, once the neuron population activity approaches the attractor trajectory, the model still fails to reproduce neuron-specific time-varying activity patterns, or the decorrelated neuronal activity observed in biological recordings of cortical neurons (Fig. 1) ^2,30,31^, neurons of the brainstem cuneate nucleus ^32^, and spinal cord neurons ^33,34^.

### How attractor trajectories and dynamic vector fields govern behavioral output

The evolving vector fields govern the spatiotemporal evolution of neural activity. Because behavior ultimately corresponds to spatiotemporal patterns of muscle activation, represented in vertebrates by the activity of the spinal motorneuron population, attractor trajectories would consequently define the behavioral output. To investigate this relationship, we implemented a highly simplified but integrated sensorimotor system consisting of a corticospinal tract (CST) input with four independent signals, qualitatively resembling the activity patterns of corticospinal neurons observed *in vivo* ^35^, a much-simplified spinal cord network, and a simple actuated biomechanical system representing the body. The sensory feedback from the biomechanical system to the spinal network was analogous to group Ia muscle spindle afferents (Fig. 4A) ^36^ and this hence formed a closed loop that included the spinal cord network.

**Fig. 4.**
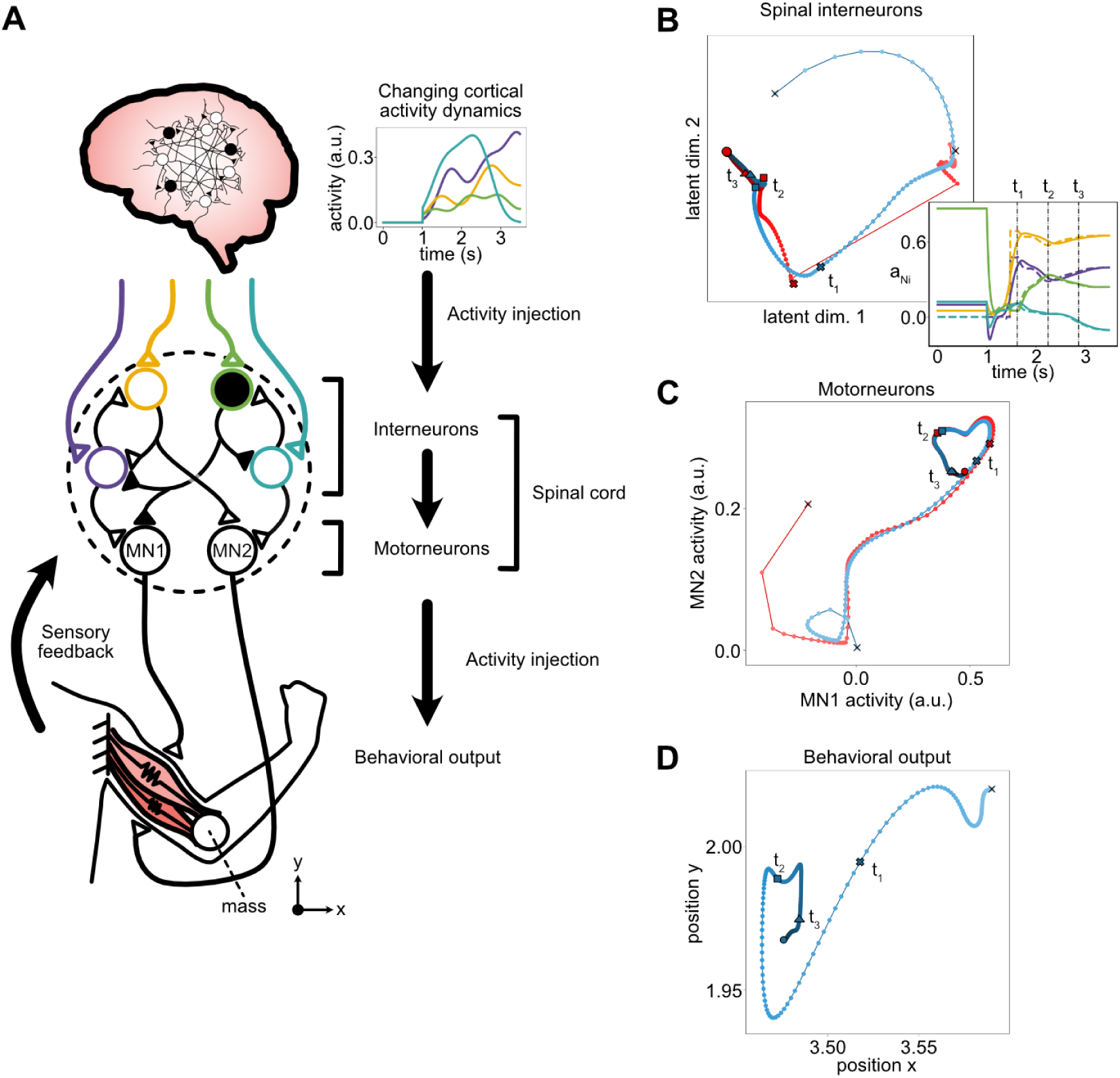
Translation from attractor trajectory to behavioral output and more decorrelated neuron activities in closed-loop through the biomechanics. **(A)** Schematic of the model used to illustrate the process from dynamic corticospinal signals to behavioral output. The cortical input drives the activity of spinal interneurons, which in turn drive the motor neuron output. Motor neuron activity ultimately determines behavioral outcomes, such as arm position. In this configuration, both spinal neurons and motorneurons integrated the sensory feedback from the body. **(B)** Projection of the attractor trajectory (red) established by the cortical-like regulatory input and the sensory feedback, that is tracked by the spinal interneuron population activity trajectory (blue). An animation is provided in Supplementary Video S1. Inset depicts interneuron activity (solid lines) as it follows the neuron-specific projection of the attractor through time (dashed lines). **(C)** The resulting interneuron activity serves as the input to the motor neuron population, establishing the attractor trajectory (red) that the motorneurons (blue) follow. **(D)** Motorneuron activity then drives the arm displacement through exciting the simulated biomechanical dynamics.

Under these conditions, the vector field of the spinal interneuron network is dynamically modified by the time-evolving inputs from the CST input and the sensory feedback to generate an attractor trajectory that controls the activity of the spinal interneuron population (Fig. 4B). Compared with the simpler systems illustrated in Figs. 2 and 3, due to the increased complexity of having multiple independent sources of external input, this integrated network produced more complex population dynamics, i.e. more diverse, decorrelated neuron activities (Fig. 4B inset). The resulting spinal interneuron activity in turn determined the attractor trajectory that drove the activity evolution of the motorneuron population (Fig. 4C), which in turn generated behavioral output through state transitions in the biomechanical system (Fig. 4D). A visualization of how the population activity continually chases the moving attractor point is provided in Supplementary Video S1.

Notably, the biomechanical properties of the biological body constitute a dynamical system. Owing to the body’s pervasive compliances, its behavior can be approximated as mutually interacting mass-spring-damper systems ^2,37^, which is analogous to the dynamics of closed-loop networks. Consequently, the body’s biomechanics can likewise be represented as a vector field ^38^, which in turn can be approximated by a neural network (Fig. 5A,B). This external dynamical system is excited by the motorneuron output, whereas the spatiotemporal patterns of the resulting sensory feedback emerge from the evolution of the dynamical state in this external system – which is constrained by the body’s anatomical structure and the physical properties of the environment. The sensory feedback, in turn, continuously impacts the vector field of the spinal network and thereby its attractor trajectories. Thus, this neuromechanical sensorimotor system ^39^ can be understood as two coupled dynamical systems that continuously impact each other’s trajectories. Within this framework, the role of the CST input is to continually bias these interacting trajectories toward a particular dynamical mode, i.e. behavior.

**Fig. 5.**
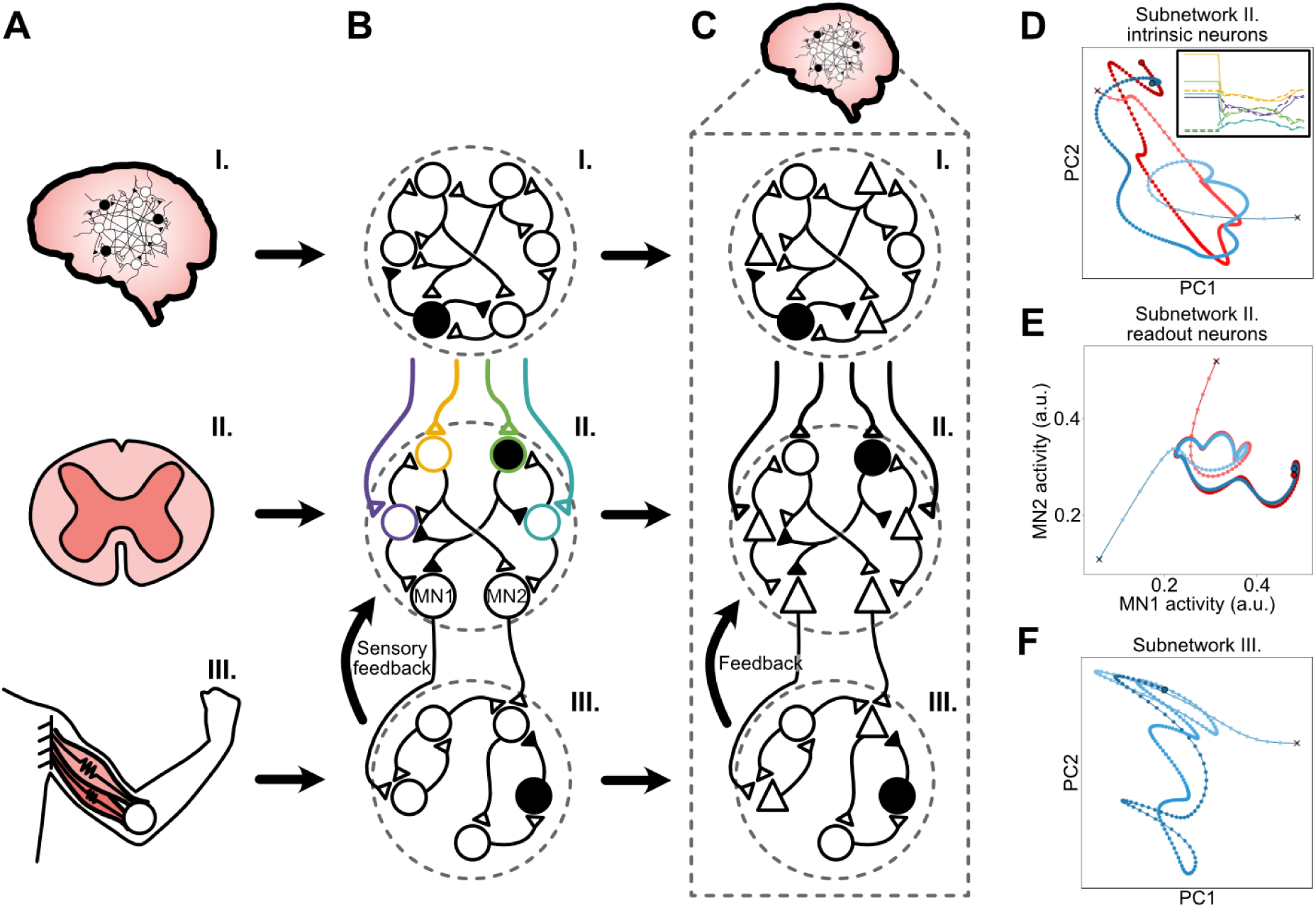
Biomechanics conceptualized as a subnetwork equivalent to the thalamocortical subnetworks. **(A)** Integrated system consisting of cortex, spinal cord, and biomechanics. **(B)** Each subsystem, including the biomechanical dynamic system, can be approximated as a neuronal network. The specific circuit implementations are highly simplified and serve only illustrative purposes to facilitate conceptual understanding. Note that the biomechanics do not represent the muscular control of the arm (the symbol is retained to facilitate comparison with Fig. 4) but instead correspond to a compartmentalized mass-spring-damper model describing the dynamics of biological skin and connective tissue ^37^ as this representation of biomechanical dynamics is more readily translated into a simple network analog. **(C)** The subnetwork architecture of the integrated system of (A) can in principle be equivalent to hypothetical circuitry structures encompassed within the thalamocortical system. Thalamic neurons (white circles) project to pyramidal cells (triangles), which in turn project back to their corresponding thalamic neurons. These local closed loops form subnetworks with their own attractor trajectories, which interact with and shape the attractor dynamics of other thalamocortical subnetworks, in an equivalent way as in the spinal neuromechanical network (roman numerals I-III refer to equivalent components across B and C). **(D-F)** Attractor (red) and activity (blue) trajectories (as in Fig 4B-D) for the two lower cortical subnetworks in (C). Note that (D) corresponds to the activity of the colored neurons in (B), whereas (E) corresponds to the lower neurons (MN1 and MN2) in the same subnetwork. The cortical-like descending command, injected by the top cortical subnetwork in (C) was identical to the input shown in Fig 4A. The combined effect of this integrated set of thalamocortical subnetwork could for example be to simulate the outcome of potential behavioral choices, i.e. imagined muscle output patterns.

This framework can also be applied to interpret the autonomous control of the neuron population activity within the cortex. The principles illustrated in Fig. 4 apply equally to intrinsic thalamocortical processing, in which closed-loop subnetworks would originate from independent thalamic neurons ^27^ (Fig. 5C). The dynamics of active conductances, prominently expressed in thalamic neurons, form an analogy to the inertial effects in the biomechanical system ^37,40^; both can self-generate dynamic activity that persists beyond, and is partly independent of, synaptic input ^6^. Since even adjacent thalamic neurons exhibit independent components of activity ^41^, each thalamic neuron tentatively forms the basis of a single thalamocortical subnetwork. Through lateral interconnections and partially shared cortical target neurons, these thalamocortical subnetworks, each with its own intrinsic vector field, will interact, in principle, in the same manner as the integrated cortico-spinal-body system (Fig. 5D-F). However, since the functional consequences of these subnetwork interactions are more readily interpreted when their outputs are expressed through a physical system, we used the biomechanical system to illustrate the remaining emergent functional principles below.

### How the same circuit can be utilized for multiple behaviors

Biological creatures can generate a wide repertoire of behaviors using the same underlying neural circuitry. In fact, the spinal cord network, for example, participates in all behaviors. Consequently, the same network needs to be capable of supporting multiple alternative attractor trajectories. We next explored how such flexibility could arise (Fig. 6). An important additional requirement is that the same behavior should remain robust to variations in corticospinal input, since a given movement can be generated by a range of different corticospinal activity patterns^20^. At the same time, the system must remain capable of transitioning between behaviors when conditions change. As detailed below, we found that both robustness and behavioral switching could emerge from perturbations in CST input, spinal interneuron initial state, or biomechanical initial state (Fig. 6).

**Fig. 6.**
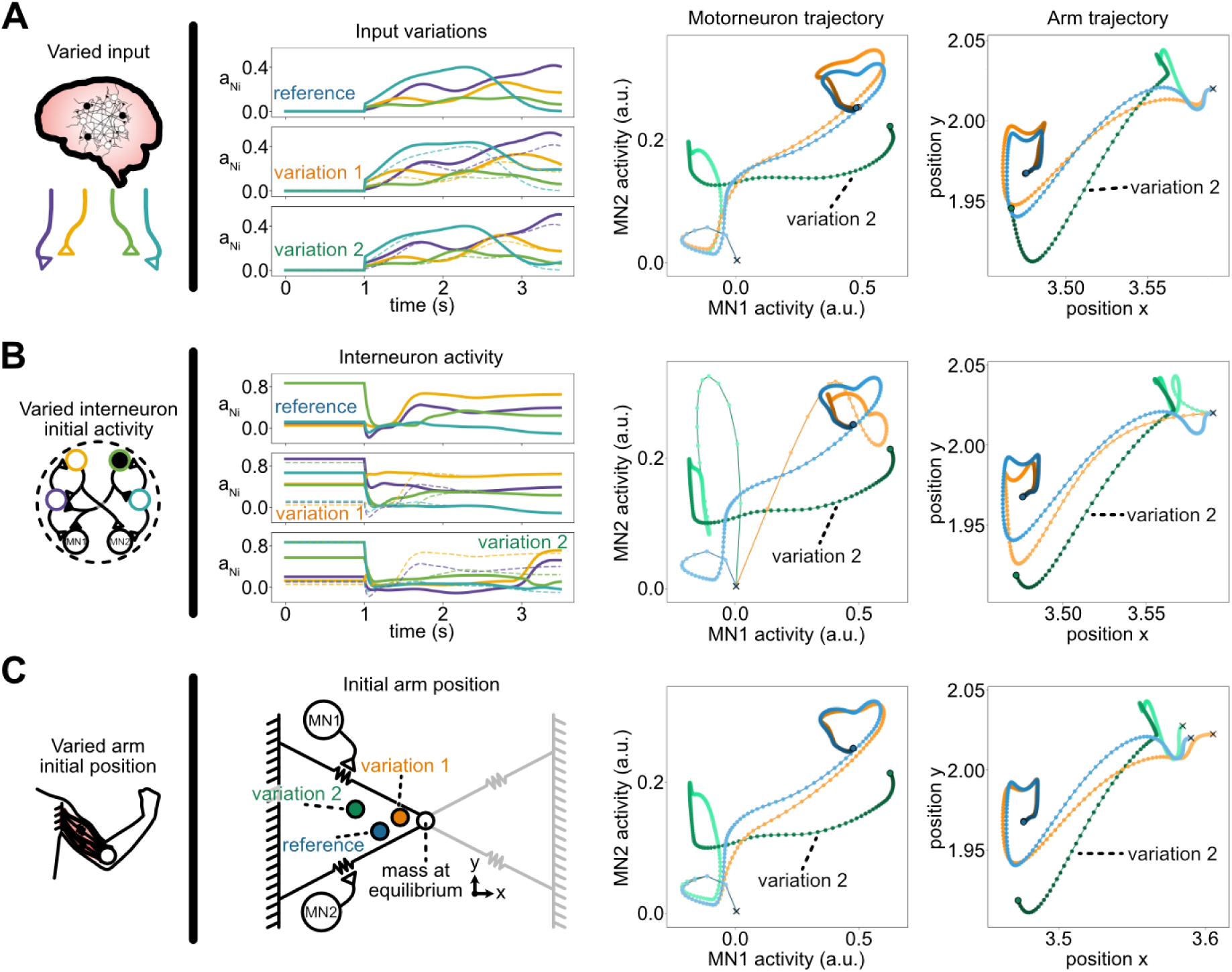
Transitions into distinct attractor trajectories and behaviors caused by critical variability across the sources of the integrated system. **(A)** Variability introduced through the descending cortical input; **(B)** the initial activity level of the spinal interneurons, and **(C)** the initial biomechanical state (arm position) at the onset of input. For each condition, a reference configuration (identical to that shown in Fig. 4) and two perturbed variants are provided. Variation 2 produces both motorneuron activity and arm trajectories that substantially differ from the other trajectories, reflecting a non-linear transition to an alternative behavior.

Within the vector field framework, a perturbation (for example activation of a sensor) can be represented as an input vector field added to the network’s intrinsic vector field. Because the input vector field is uniform across the activity state space, whereas the network’s vector field is spatially structured, the effect of a given perturbation depends on the network state at which it is received. The same perturbation can have a greater influence when the network state lies in a region where the intrinsic dynamics evolve comparatively slowly (i.e., where the intrinsic vector lengths are small), whereas its effect could be reduced in regions where the dynamics are faster. Beyond the immediate state in which the input is received, the local structure of the network’s intrinsic vector field also determines the perturbation’s long-term effects. When a perturbation is sufficiently small relative to local dynamics, it produces only a transient deviation in the activity trajectory, which we refer to as a “wobble”, before the trajectory returns toward its original course. In contrast, a larger perturbation may move the system into the basin of attraction of a different attractor trajectory, resulting in a qualitatively different behavioral outcome. This distinction is illustrated by two variations in CST input, one producing a wobble and the other redirecting the integrated dynamic system into an alternative trajectory (Fig. 6A), i.e. behavior.

Variability in the example integrated system can arise from multiple sources. Variability in corticospinal input may reflect intentional modulation or spontaneous neural fluctuations. A second source of variability in the model was the initial state of the spinal interneuron population. Different initial activity distributions corresponded to different starting points in state space, allowing identical corticospinal inputs to produce different outcomes and, in some cases, distinct behaviors (Fig. 6B). This finding highlights the importance of the cortex inducing preparatory activity in the spinal interneuron network before movement execution ^42^.

Behavioral variability also arose from differences in the arm’s initial biomechanical state (Fig. 6C). Each initial position corresponded to a specific distribution of muscle tension across the two muscles. Altering this initial state of the mechanics caused the spatiotemporal sensory feedback patterns to the spinal circuitry to change. Hence, the changes in pre-tension biased the system toward different attractor trajectories and, therefore, behavioral outcomes, despite identical states in the corticospinal input and the spinal network. Together, these results demonstrate that the attractor trajectory-based framework accounts for how behavioral variability can arise from specific changes in CST inputs, spinal network states, or biomechanical conditions, while other perturbations instead resulted in unchanged behavior, reflecting also a behavioral robustness of the system. Fig. S3 illustrates that the same principles emerged when the sensory feedback was provided solely to the spinal interneurons rather than also including the motorneurons as projection targets. These findings highlight the importance of preparatory activity to align the states of all the different dynamical systems in the interaction, thereby facilitating the behavioral performance.

## DISCUSSION

We addressed the emergent functionality from the nested closed loop network dynamics of the brain and how it despite its inherent complexity can be understood through the vector field framework. We argue that this perspective provides a new angle to understand the foundations of brain operation and how it emerges from basic neural principles. Membrane capacitance and channel leak introduce an attractor point into the vector fields (Fig. 2). Since the vector field of a network is fixed, its activity will converge to that attractor point. But when the network is mutually interconnected with another dynamical system, the network’s vector field structure dynamically evolves (Figs. 3-6) which gives rise to decorrelated spatiotemporal neural activity patterns qualitatively similar to those observed *in vivo* (Fig. 1). Whereas dynamic sensory inputs can only arise from the body biomechanics when it is excited, time-continuous vector field changes are also caused by neuron-intrinsic active conductances that self-generate dynamic activity even without synaptic, e.g. sensory, inputs. Independent self-generated activity across thalamic neurons will continually reshape the thalamocortical vector field, thereby producing internally generated cortical activity trajectories (Figs 1,3,5). This provides a mechanistic understanding of the essence of ‘thoughts’ as a process which is needed to anticipatively time state initiation in body-world interactions (Fig. 6) based on acquired interpretations of cumulative sensory cues, rather than merely immediately responding to such sensory stimuli ^1^.

### Relationship to the literature

The attractor trajectories described here differ from the attractor frameworks commonly applied in neuroscience literature. One prominent example is discrete attractors, which are fixed points in the network’s state space toward which the system flows and eventually settles. Such attractors have been proposed as a mechanism for memory representation ^43,44^. A closely related framework is that of continuous attractors ^45,46^, which consist of a continuous set of fixed points that collectively form a low-dimensional structure, often referred to as a manifold. Along this manifold, there is no genuine movement, whereas perturbations in directions perpendicular to the manifold are driven back toward it. Another framework is the limit cycle ^47,48^, in which the system’s dynamics evolve continuously along a closed, periodic trajectory. In contrast, the attractor trajectories described here resemble limit cycles in that activity continuously flows along the attractor, but they are fundamentally non-periodic. Rather than repeatedly traversing a closed orbit, neural activity progresses along an open trajectory through the state space, making these attractors more appropriately conceptualized as directed flows within the network’s vector field.

### How multiple diverse behaviors can be supported by the same network

The complexity arising from nested closed loops provides an evolutionary advantage by enabling a greater diversity of behavior (see ^13,19^). We showed that diverse behavior can emerge through interactions between two dynamical systems, at least one of which possesses multiple solutions. These solutions may reside within the neural network itself, arising from its connectivity structure (Figs. 4,6), or from factors external to the network, originating in the dynamical systems with which it interacts. The biomechanics of the body provide an example of such an external dynamical system, where multiple mechanically stable solutions have been identified ^49^. The series of coupled mechanical effects within each such solution gives rise to sensory dependencies with specific spatiotemporal structure^2^. The sensory dependencies in turn give rise to a specifically evolving impact on the vector field, inducing a specific attractor trajectory in the downstream subnetwork, in the present paper illustrated by the spinal cord. Because attractor trajectory selection depends on the initial state of both the spinal network and the mechanical system (Fig. 6), our results indicate that the cortex preparatively needs to ensure appropriate initial states before movement onset, which is a possible explanation for the evolutionary origin of anticipatory cortical control.

When there are multiple attractor trajectories in the neuromechanical system, perturbations can induce transitions between them. The effect of a given perturbation, of sensory or (thalamo-)cortical origin, depends on the geometry of the vector field within the current part of the activity state space. As shown in Fig. 6, perturbations may produce either transient deviations (wobbles) that decay back to the original attractor trajectory, or sustained cascades of activity that drive the system into a different trajectory, depending on the positions within the network and the biomechanics state spaces when the perturbation occurs. State space regions of low susceptibility give rise to activity trajectories that are robust to external influences, whereas regions of high transition susceptibility facilitate switching between attractor trajectories. We refer to this regime as operating near criticality, where small perturbations can produce large behavioral changes. Criticality allows flexible and adaptive behavioral transitions to be triggered by a low effort in terms of synaptic inputs. The enhanced responsiveness afforded by such criticality could greatly facilitate learning by enabling the network to more readily acquire and retain multiple functional solutions.

### Alignment in the dynamics domain - a possible role for learning across the CNS

The closed-loop coupling between the neural and biomechanical systems, in which the output of each system serves as the input to the other, can produce mutually reinforcing dynamics that make behavior more automated, smoother and less reliant on corrections. When the neural and biomechanical trajectories evolve within their respective vector fields such that their mutual interactions reinforce or sustain each other’s trajectories, rather than causing them to fizzle, the two systems can be said to be well aligned. Specifically, the initial biomechanical state, together with neural excitation, generates a sequence of biomechanical state transitions that in turn produce corresponding sensory state transitions. These sensory transitions reshape the vector field, and thereby the attractor trajectory, of the spinal network, defining the spatiotemporal pattern of motor neuron activity (Fig. 6). The closer the motor output is aligned with the ongoing biomechanical state trajectory, the more effectively the mutual excitation between neural and biomechanical dynamics is reinforced, which would result in smoother and more stable movement. During such aligned dynamics, unintended neural and sensory fluctuations, as well as spike firing noise ^50,51^, are then largely expressed as wobbles rather than sustained deviations. In contrast, when the neural vector field is poorly aligned to the biomechanical vector field, the resulting trajectory misalignment ^52^ would increase the demand for corrective control and could cause the behavior to fail altogether.

In our example system, cortical input and sensory feedback constituted the primary sources of independent input to the spinal circuitry. However, the CNS also contains numerous reactive brainstem circuits, including vestibular, reticular, and other descending systems. These circuits receive independent sensory inputs (e.g. vestibular inputs) and exert their corrective action through closed-loop control, where their impact on motor output feeds back through the body to impact the sensory inputs that drive them ^1,53^. Hence, they form additional, largely independent subnetworks that impact the spinal vector fields. Such an organization of interacting subnetworks is also formed by closed-loop thalamocortical circuits ^27^ (see Results text) that form a system of interacting dynamical subnetworks (Fig. 5C-F). Self-generated thalamic neuron activity would provide for sensory-independent processing, which is needed for anticipative state alignment (Fig. 6) and to imagine future states for a simulated behavioral choice^1^. Competing corrective actions among the nested dynamical systems across the CNS, and to anticipate them, is a mechanistic basis for reinterpreting neural population activity ^19^.

In this conceptual framework, a primary objective of learning would be to align the structure of the spinal neural network with the dynamics of the body’s biomechanics. Achieving this requires the network to adapt its connectivity, i.e. its synaptic weights, such that its dynamics mirror the structure of the biomechanical dynamics ^2,52^. Anticipatory cortical function (Fig. 6) implies a need to learn how to integrate a preceding history of cross-modal sensory dynamics into the nested recurrent dynamics of interacting cortical subnetworks so that they, independently of ongoing movement activity in the spinal cord, anticipatively act to prevent reactive corrective actions arising during specific body-world interactions ^1^. Hence, rather than viewing behavior as the outcome of non-constrained neural computation, this framework implies that behavior emerges from the alignment of mutual dynamic interactions between neural subnetworks and biomechanics. Learning would align those interaction dynamics to maintain stable, smooth trajectories while also allowing for criticality effects that would support on-the-fly adaptability during more uncertain behavioral phases and during perturbation.

### Limitations

We deliberately kept the modelled networks as small and generic as possible, to focus on explaining the ‘atomic’ functional consequences of the conceptual framework, rather than championing any specific features through the model implementation. Hence, the possible specific roles of more specific neuronal subtypes, other than the distinction between excitatory and inhibitory neurons, were not explicitly addressed here. But it should be noted that specific neuron subtypes identified in the literature would in the fast computational domain we considered here mainly differ in regard to their specific intrinsic activation function and the duration of the synaptic influence it is exerting on its postsynaptic targets, which are implementable in this framework (see Ref. 15). Furthermore, our networks were very small compared to a real brain. But as we have previously found, the computational architecture naturally does not change just because the neuron population grows larger ^2^. However, the simple structure of our simulated dynamic biomechanical system, with very few components and limited nonlinear behavior, limited the full range of the complexity of the behavior in the neural-biomechanical system compared to any relevant biological system (vertebrates) in comparison.

## MATERIALS AND METHODS

### In vivo neocortical recordings

*In vivo* cortical recordings were obtained from previously published datasets in anesthetized rats^13,54^ and awake, head-fixed mice^55^. All recordings were acquired using Neuropixels multi-electrode arrays^55^ (384 recording channels).

For the anesthetized recordings, rats were anesthetized with ketamine-xylazine. From that dataset, we used previously sorted and manually curated cortical units^13,54^. Probes were inserted orhogonally to the cortical surface, and only units recorded at depths <1.7 mm (maximum cortical depth in rats) were included. Spike sorting was performed using Kilosort2.5^56^. Two recordings were analyzed: one from the primary somatosensory cortex (S1) and one from the primary visual cortex (V1).

Awake recordings were analyzed from the International Brain Laboratory (IBL) Brain-Wide Map dataset^57^, available through DANDI Archive (Dandiset 000409, version 0.260309.1324). Recordings were obtained using Neuropixels 1.0 (probe type 3B2) probes, and analyses were restricted to units residing on channels that were anatomically assigned to neocortical regions according to the Allen Common Coordinate Framework. The complete list of recording sessions included in the analysis is provided in Supplementary Table 1. Spike sorting was performed using Kilosort4^56^ on the Neuropixels recordings, and only units classified as “good” by the sorting pipeline were retained.

For both datasets, units were further filtered based on their recording presence. The recording was divided into 1-second bins, and a unit was considered present in a given bin if at least one spike occurred within that bin. Only units present in at least 80% of the recording bins were retained. This filtering resulted in 128, 27, 25, and 208 sorted units from the anesthetized rat S1, anesthetized rat V1, and the two awake mouse recordings, respectively.

For qualitative evaluation of population dynamics, spiking activity from the V1 recording was convolved into a time-continuous representation. Spike trains were first binned in 10 ms intervals and subsequently smoothed with a 50 ms Gaussian kernel to estimate instantaneous firing rates. Population activity was then projected into the first three dimensions obtained using both linear (PCA) and nonlinear (UMAP) dimensionality reduction techniques, enabling visualization of the resulting population dynamics.

### Neural stationarity measure

Stationarity was assessed at both the single-neuron and the neuron population levels. Individual neurons were considered stationary when they exhibited low variability in their interspike intervals (ISIs). Firing variability was quantified using the coefficient of variation (CV), defined as the ratio of the standard deviation (ro) to the mean (mu) of consecutive ISIs (Eq. 1). A CV of zero indicates perfectly regular firing, while higher values correspond to greater variability.

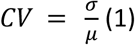

To characterize changes in firing variability over the course of a recording, CV was evaluated for overlapping ISI sequences of varying lengths. Specifically, CV values were computed for sequences containing 10, 20, …, 100 consecutive ISIs. For each sequence length, a sliding-window approach was used in which successive windows overlapped by 50%. Each CV estimate was associated with the time interval spanned by the corresponding ISI sequence, allowing variability to be examined across both short and long temporal scales. This procedure provided a time-resolved measure of neuronal firing variability throughout the recording for individual neurons.

After computing CV values for individual neurons, we evaluated the extent to which stationarity occurred at the population level. Following the criterion of ^26^, ISI sequences with CV < 0.6 were classified as exhibiting low variability and therefore considered stationary. Population-level stationarity was defined as periods during which stationary intervals overlapped across multiple neurons. For each time period, we quantified both its duration and the proportion of the recorded population that exhibited stationary activity.

### Conductance-based non-spiking neuron model

To investigate neuron population dynamics, we built network models using a conductance-based, non-spiking neuron model first introduced in ^28^ (also described in ^15^). This model, referred to as the Linear Summation Neuron Model (LSM), encompasses a linear summation of synaptic input that then contributes to the neuron’s dynamic state transition. The activity level of the post-synaptic neuron depends on the presynaptic input(s) and can be summarized by

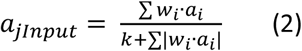

where the summation iterates over all presynaptic neurons *i* of neuron *j*. The activity level of presynaptic neuron *i* is indicated by *a_i*, and the weight (or strength) of the connection from neuron *i* to neuron *j* is indicated by *w_i*. The denominator includes a factor that divides the incoming presynaptic inputs by the total currently active presynaptic activity, hence producing a shunting effect that scales with the total synaptic activity. The *k* parameter acts as a form of normalization which aims to reflect the constituent leak channels that give rise to the neuron’s membrane potential. Since the membrane surface density of leak channels is roughly consistent across the dendrites ^58,59^ and the membrane area scales with the number of synapses the neuron receives, the *k* parameter is scaled with the number of synapses.

The dynamics of the neuron model is described as

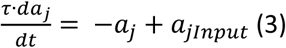

where τ is the neuron membrane time constant, which is referred to as the dynamic leak in ^28^, and *a_jInput* is obtained from Equation 1. In a biological cell, the time constant depends on both the capacitance and the resistance (leak) of the membrane, and because of the capacitive effect, the membrane time constant can be said to constitute a type of state memory in the neuron. It should be noted that in this instantiation of the LSM model, a neuron’s resting membrane potential corresponds to an activity level of zero, and the dynamic leak will over time tend to bring the neuron back towards its resting state. Whenever the activity level of the LSM neuron exceeds the resting state value, it provides synaptic input to its downstream neurons.

This neuron model represents the rate code of neuronal output as a proxy for underlying activity levels. This approach is consistent with many studies aimed at characterizing neuron population interactions ^60–62^. In biophysical terms, each presynaptic spike induces a synaptic potential in the postsynaptic neuron, which in turn influences its output activity. Although a spike in a presynaptic neuron may last for less than 1 ms, the synaptic potential that it induces in the postsynaptic neuron typically persists for many tens of milliseconds ^63,64^, gradually decaying over time. When the interspike interval of the postsynaptic neuron is shorter than the duration of the synaptic potential, successive postsynaptic potentials overlap and summate in a time-dependent manner. This temporal integration effectively gives rise to a rate-based representation, consistent with the modeling framework used here.

### Vector field framework

To illustrate the principles of network function in an integrated system, we employed an extension of the vector field framework introduced in ^15^. In this framework, a neuronal network is represented as a vector field whose dimensionality equals the number of neurons in the network, with each dimension corresponding to the normalized activity level of a single neuron. The activity distribution over the neuron population uniquely defines a location, or state, within this n-dimensional space. The space encapsulating the totality of possible activity states can therefore be referred to as the state space of the network. Neuronal activity evolves through synaptic interactions, where neurons influence one another according to their connectivity and activation dynamics. The effect of these interactions can be expressed as a vector that describes the instantaneous change in network state. Because this vector depends on both the current activity levels of the neurons and the synaptic weights connecting them, evaluating it across all possible network states yields a vector field.

To provide an intuitive understanding of the core principles, we first considered a simple two-neuron network composed of mutually interconnected excitatory neurons. The low dimensionality of this system allows the full-dimensional vector field to be visualized directly. Although all example networks used in this study were small and used the LSM as its activation function, the underlying principles are general and can be scaled to larger networks with arbitrary connectivity patterns and activation functions.

### Simplified model of an integrated system

To illustrate the core operating principles of an integrated system, we implemented a system of interacting networks representing a highly simplified model of cortical control of spinal cord circuitry for behavior generation through a biomechanical model. The corticospinal pathway was modeled as four independent channels that projected activity to a reduced network representing the spinal cord. This simplified spinal circuit consisted of four interneurons, each receiving input from one cortical channel, and two motorneurons. The interneurons were connected to one another and projected to the two motorneurons that serve the functional role of output neurons of the neural system. The motorneurons then actuated the biomechanical model (described below), closing the sensorimotor loop through sensory afferent feedback to the spinal circuitry.

### Biomechanical model

To assess the behavioral effects of our simulated integrated network, we developed a simple biomechanical arm model whose muscle-like actuators were driven by motorneurons in the spinal circuit. The behavior of the integrated neuromechanical system was measured as the trajectory of the arm in response to neural activation from the cortex. The arm model consisted of two muscles, innervated by one motorneuron each, operating in partial opposition to each other. Sensory feedback from these muscles was provided to both motorneurons and spinal interneurons.

The arm was situated in a two-dimensional environment, with one end of each muscle anchored to a fixed wall and the other attached to a moveable mass. The muscles’ anchor points were positioned vertically one unit length apart. Activation of the motorneurons induced muscle contraction proportional to the level of neural activity. As motorneuron activity decreased or ceased, the arm returned toward a state of muscle extension through the action of two passive springs, one associated with each muscle, which opposed the contractile forces. This could be thought of as gravity acting on the mass position when the muscles relaxed. The resulting two-dimensional trajectory of the mass in response to muscle activation was defined as the behavioral output.

The equations governing the manipulation of the mass position and the proprioceptive sensory feedback dynamics were adapted from ^36^. The force exerted by each muscle depended both on the muscle’s activation (i.e. the motorneuron output) and its length relative to its maximal length. This force is further impacted by the muscle’s viscous properties, which provide a damping effect. The muscle dynamics are described in Eq. 3, where A represents muscle activation, L denotes the maximal length, and D is the viscosity-related damping coefficient. Subscripts i indicate the specific muscle (either 1 or 2), with x_i_ and xdot_i_ referring to the muscle’s length and velocity, respectively. A complete list of variables and their corresponding values is provided in table S2.

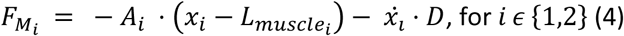

The force exerted by the counteracting passive springs is given by Eq. 5, where k represents the stiffness, and subscripts (either 1 or 2) denote the specific passive spring.

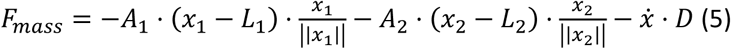

The total force acting on the mass, resulting from the combined effects of the two actuated muscles and their opposing passive springs, is expressed in Eq. 6.

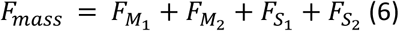

Proprioceptive sensory feedback to the spinal cord was modeled using group Ia muscle spindle afferents, which capture a property also shared by group II muscle spindles ^36^. The dynamics of these spindles are described in Eq 7 and Eq 8, where L and V denote length and velocity, respectively. In this study, only Ia afferents were included as sensory input (whereas a component of the group II sensor activation is a factor in calculating the Ia afferent activity, see Eqs 6 and 7). Their outputs were additionally subjected to thresholding at different activity levels ^65^.

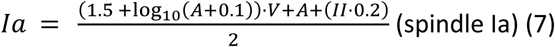

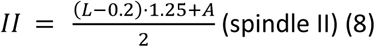

It is important to note that while motorneurons activity could theoretically be negative (e.g. during predominantly inhibitory presynaptic input), the output activity of the motorneurons cannot fall below zero. This ensures that muscle activation is only modulated by non-negative motorneuron output.

## Funding

EU H2020 FETOpen project# 829168 ‘ph-coding’

VINNOVA

Swedish Medical Research Council

## Author contributions

Conceptualization: SS, HJ

Methodology: SS, HJ

Investigation: SS, HJ

Visualization: SS, HJ

Funding acquisition: HJ

Project administration: HJ

Supervision: SS, HJ

Writing – original draft: SS, HJ

Writing – review & editing: SS, HJ

## Data, code, and materials availability

The data and code that support the findings of this study will be publicly available upon publication.

## Supplementary Materials

Figs. S1 to S3

Supplementary Video S1

Tables S1 and S2

