## Supplementary Figures for "How brain operation emerges from its nested recurrent circuitry extending into the body"

**The PDF file includes:**

Figs. S1 to S3
Caption for Supplementary Video S1
Tables S1 and S2

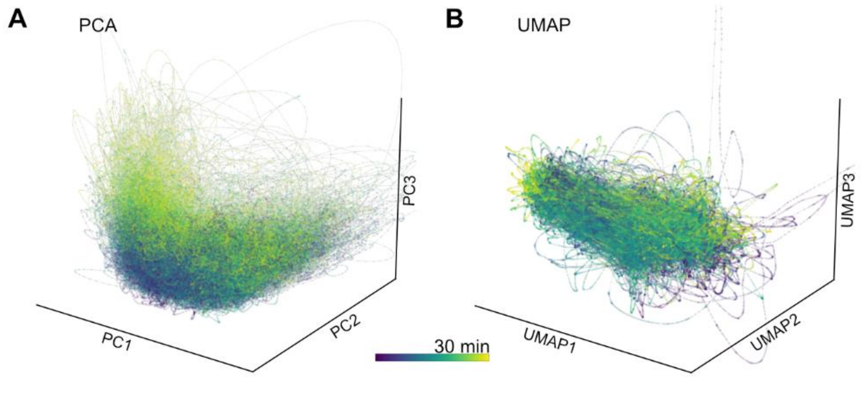


**Fig. S1.**

**Representations of the dynamically changing cortical population activity in the awake mouse**. Similar display as in Fig. 1B,C. Note that in the end of the 0-30 mins recording period, the population activity enters a new part of the PC space (panel A), which suggests the presence of some drift in the unit sorting.

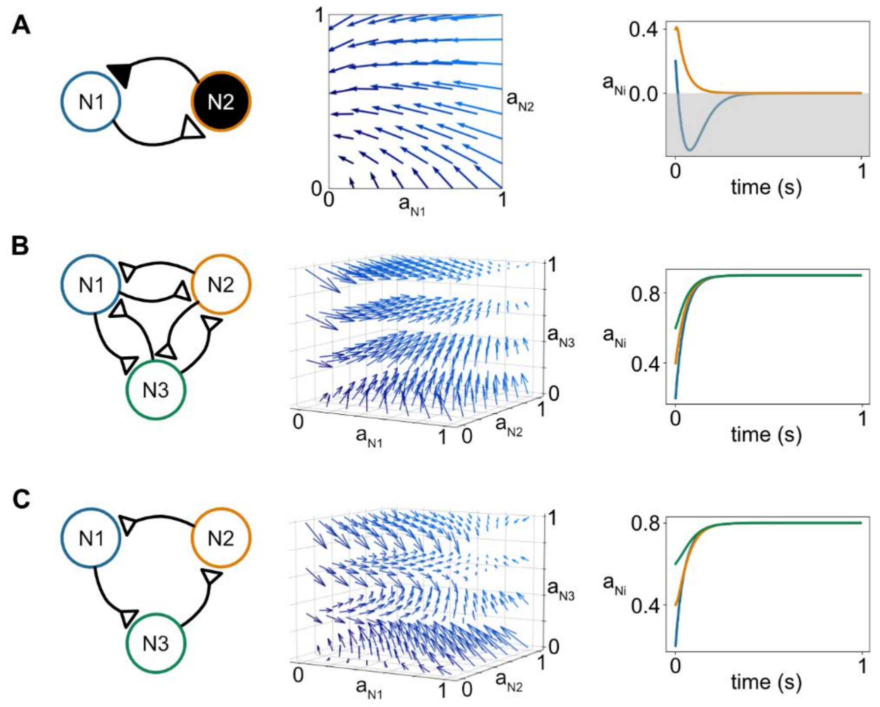


**Fig. S2.**

**The vector field representation can be extended to arbitrary network configurations.** (A) Introducing inhibitory neurons reduces the overall excitation in the network. In the example recurrent excitatory-inhibitory network (left), the specific weight combination leads to a removal of the non-zero attractor point (middle). The attractor point now occurs at zero activity for both neurons, as confirmed by a simulation (right), where activity converges to and remains at this state. (B) Same network as in Fig. 2 but extended with one more neuron (fully connected network). The three-dimensional vector field reveals its non-zero attractor point, to which the activity of the neuron population converges (right). (C) In a more sparsely connected network (left), the structure of the resulting vector field (middle) differs from that of the fully connected case. Reduced excitatory connectivity lowers the overall network excitation, shifting the attractor point to a lower, but still non-zero, activity level (right). Note that neither conduction delays nor implementing the neurons as spiking neurons would alter the principle of activity convergence towards the attractor point that emerges across these examples, but the probability of reaching the non-zero attractor point could alter.

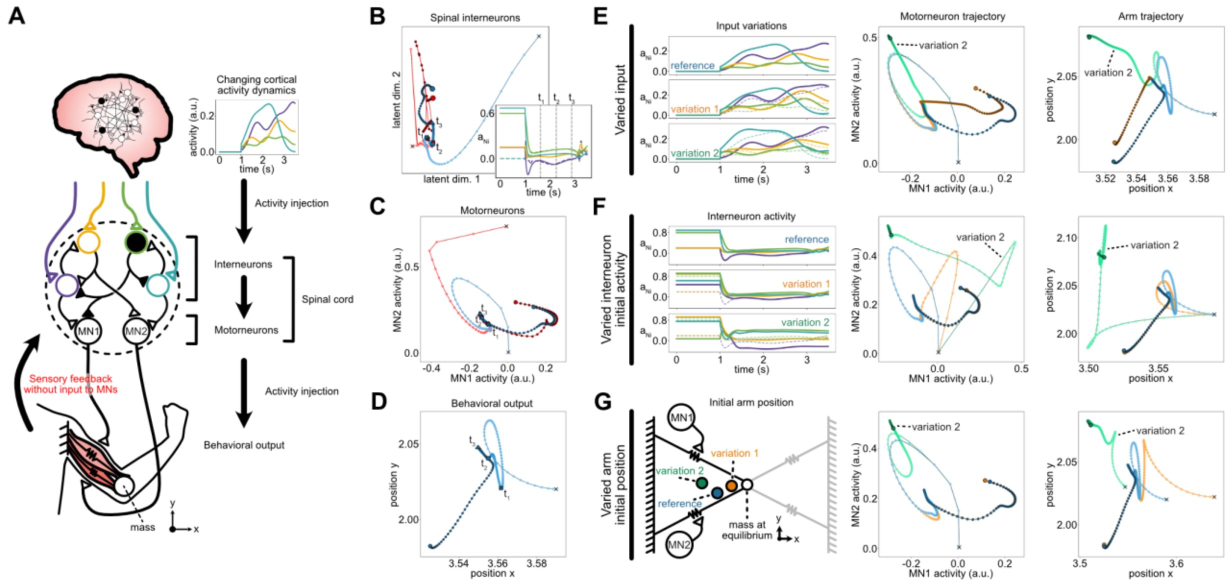


**Fig. S3.**

**Attractor trajectories and multiple behavioral output without direct sensory feedback to motorneurons**. (A) Schematic of the modified integrated system shown in Fig. 4A. The key difference lies in the sensory afferent pathways, where feedback projections only innervate spinal interneurons. The spinal circuitry, motorneuron innervation of muscles, and biomechanics remained unchanged. The magnitude of the cortical-like descending input was reduced to balance its influence on network dynamics with the overall reduction in network excitation caused by the removal of motorneuron sensory afferents. (B-D) Spinal interneuron, motorneuron, and behavioral output, respectively, as in Fig. 4B-D, for the modified system configuration. (E-G) The effects of variability are analogous to those shown in Fig. 6A-C, with differences limited to the exact trajectories resulting from the altered feedback connectivity.

**Supplementary Video S1**

Illustration of the attractor trajectory (red) and the travel through its activity state space (blue) of the network illustrated in Figure 4.

**Table S1.**
Awake recordings used from the IBL dataset.

| Subject | IBL eid | Probe | Device | Channels | Num neurons | DANDI asset |
|---|---|---|---|---|---|---|
| ZM-2241 | ff4187b5-4176-4e39-8894-53a24b7cf36b | 3B2 | Probe00 | 220-384 | 25 | sub-ZM-2241_ses-ff4187b5-4176-4e39-8894-53a24b7cf36b_desc-raw_ecephys.nwb |
| UCLA030 | 7f5df7eb-cf36-4589-a20a-14b535441142 | 3B2 | Probe00 | 1-285 | 208 | sub-UCLA030_ses-7f5df7eb-cf36-4589-a20a-14b535441142_desc-raw_ecephys |

**Table S2.**

Parameter definitions and values.

| Parameter | Symbol | Value |
|---|---|---|
| Muscle activation | A | [0,1] |
| Muscle maximal length | L_muscle | 3.0 |
| Damping | D | 0.246 |
| Mass | M | 0.01 |
| Stiffness | K | 3 |
| Passive spring length | L_spring | 2.0 |
